# Maximizing ecological and evolutionary insight from bisulfite sequencing data sets

**DOI:** 10.1101/091488

**Authors:** Amanda J. Lea, Tauras P. Vilgalys, Paul A.P. Durst, Jenny Tung

**Affiliations:** Department of Biology, Duke University, Box 90338, Durham, NC 27708, USA; Department of Evolutionary Anthropology, Box 90383, Durham, NC 27708, USA; Department of Biology, University of North Carolina at Chapel Hill, CB #3280, Coker Hall, Chapel Hill, NC 27599; Institute of Primate Research, National Museums of Kenya, P. O. Box 24481, Karen 00502, Nairobi, Kenya; Duke University Population Research Institute, Box 90420, Durham, NC 27708, USA

**Author notes:** Other author.

**Keywords:** DNA methylation, bisulfite sequencing, cell type heterogeneity, population structure, mixed effects models, ecological epigenetics

## Abstract

The role of DNA methylation in development, divergence, and the response to environmental stimuli is of substantial interest in ecology and evolutionary biology. Measuring genome-wide DNA methylation is increasingly feasible using sodium bisulfite sequencing. Here, we analyze simulated and published data sets to demonstrate how effect size, kinship/population structure, taxonomic differences, and cell type heterogeneity influence the power to detect differential methylation in bisulfite sequencing data sets. Our results reveal that the effect sizes typical of evolutionary and ecological studies are modest, and will thus require data sets larger than those currently in common use. Additionally, our findings emphasize that statistical approaches that ignore the properties of bisulfite sequencing data (e.g., its count-based nature) or key sources of variance in natural populations (e.g., population structure or cell type heterogeneity) often produce false negatives or false positives, thus leading to incorrect biological conclusions. Finally, we provide recommendations for handling common issues that arise in bisulfite sequencing analyses and a freely available R Shiny application for simulating and performing power analyses on bisulfite sequencing data. This app, available at www.tung-lab.org/protocols-and-software.html, allows users to explore the effects of sequencing depth, sample size, population structure, and expected effect size, tailored to their own system.

## Introduction

DNA methylation – the covalent addition of methyl groups to cytosine bases – is a gene regulatory mechanism of well-established importance in development, disease, and the response to environmental conditions^1–3^. In addition, shifts in DNA methylation are thought to 40 contribute to the speciation process and the evolution of trait differences between taxa^4–6^, in support of the idea that gene regulation plays a key role in evolutionary change. Because of its contribution to phenotypic diversity, interest in DNA methylation from the ecology and evolutionary biology communities is high^7–10^. This interest has been further encouraged by the development of sodium bisulfite sequencing, a cost-effective approach that allows researchers to measure genome-wide DNA methylation levels at base-pair resolution in essentially any organism^11–13^.

Such approaches rely on sodium bisulfite treatment of DNA followed by high-throughput sequencing, producing data sets that are collectively termed “bisulfite sequencing data.” These data have properties (discussed in the following section) that differ in key ways from other common types of sequencing-based functional genomic data, such as RNA-seq data. Consequently, several statistical approaches have been developed that are specifically tailored to bisulfite sequencing data sets^14–17^ (Box 1). However, the development, application, and evaluation of these tools has primarily focused on biomedical questions or model systems, with an emphasis on case-control studies and experimental manipulations in a restricted set of species^18–20^. In contrast, ecologists and evolutionary biologists often study non-model organisms, environmental gradients that do not follow a case-control design, and natural populations characterized by complex kin or population structure. They are also typically more limited in their ability to sample pure cell types, and may be interested in effects that are smaller than those reported in the context of major perturbations like cancer or pathogen infection^21–23^. Notably, all of these properties can affect statistical power.

Our goal in this review is to outline considerations for the analysis of bisulfite sequencing data, tailored specifically to concerns that commonly arise in ecological and evolutionary studies. We first discuss how high-throughput bisulfite sequencing data sets are generated, and how this process leads to several idiosyncrasies that must be taken into account in data analysis. Next, we consider four properties common to ecological and evolutionary data sets that can influence power to detect differential methylation: moderate effect sizes, kinship/population structure, taxonomic differences in DNA methylation patterns, and cell type heterogeneity. We analyze both simulated and published empirical data sets to demonstrate how these four features can affect the power and biological interpretation of differential methylation analysis. Finally, we provide recommendations for handling each issue, with the aim of facilitating robust, well-powered studies of DNA methylation’s role in ecological and evolutionary processes.

## Common properties of high-throughput bisulfite sequencing data sets

High-throughput bisulfite sequencing protocols, such as whole genome bisulfite sequencing (WGBS^13^) or reduced representation bisulfite sequencing (RRBS^11^), rely on the differential sensitivity of methylated versus unmethylated cytosines to the chemical sodium bisulfite (Figure 1). Specifically, treatment of DNA with sodium bisulfite converts unmethylated cytosines to uracil (replicated as thymine after PCR) but leaves methylated cytosines unchanged (in vertebrates, most DNA methylation occurs at cytosines in CG motifs, while, in other taxa, cytosines in CHG and CHH are also commonly methylated^24,25^). DNA methylation level estimates at a given site can thus be obtained via high-throughput sequencing of bisulfite converted DNA, by comparing the relative count of reads contain a cytosine (C), which reflect an originally methylated DNA base, to the count of reads containing a thymine (T), which reflect an originally unmethylated version of the same base. Current bisulfite sequencing protocols require low amounts of input DNA^26,27^, avoid the use of species-specific array platforms, and can be applied to organisms without a reference genome^28^, making them an increasingly popular choice for ecologists and evolutionary biologists^29^.

**Figure 1.**
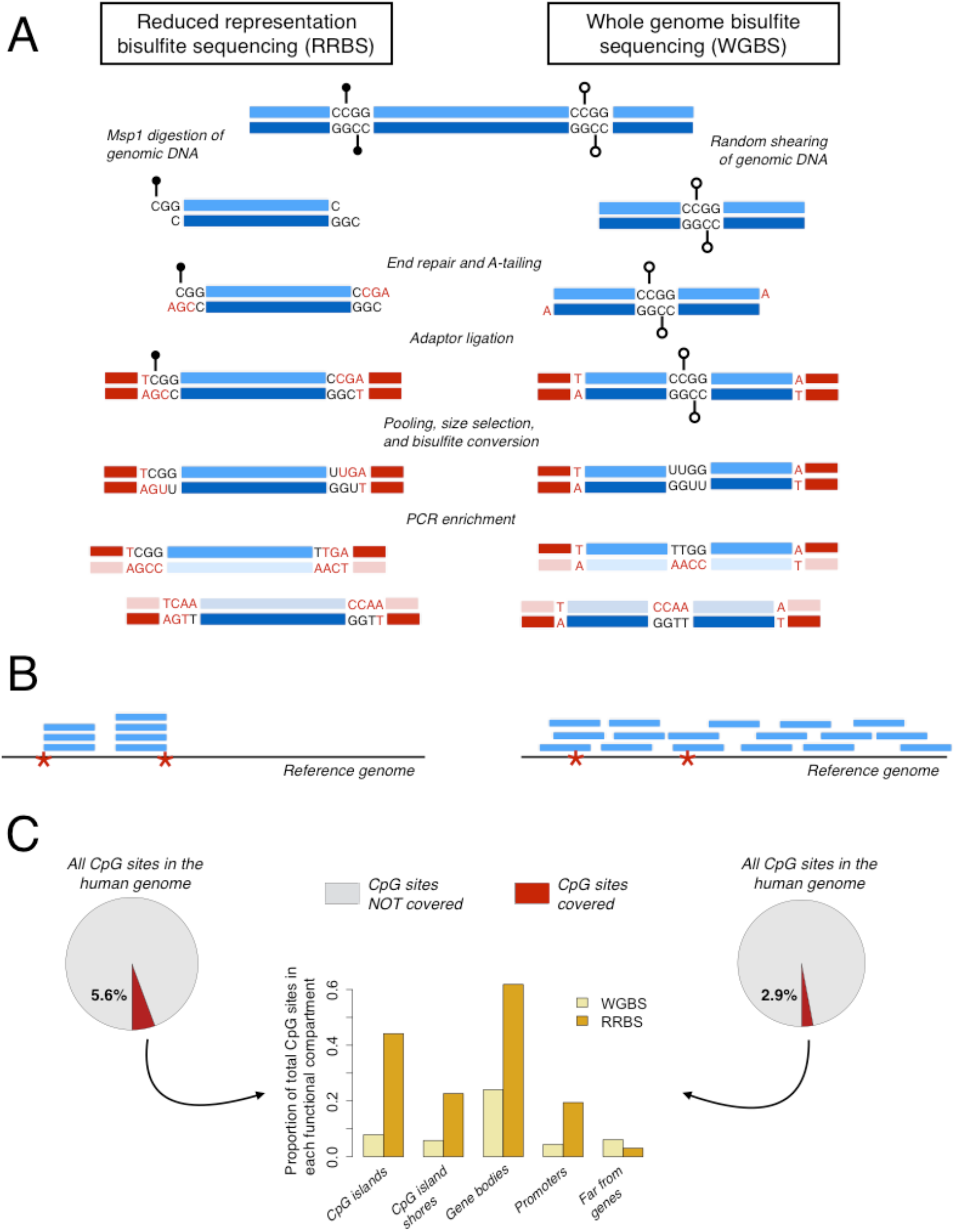
Overview of reduced representation bisulfite sequencing (RRBS; left side of figure) and whole genome bisulfite sequencing (WGBS; right side of figure). (A) Steps required to prepare an RRBS or WGBS library from genomic DNA. Methylated CpG sites are denoted with black lollipops and unmethylated CpG sites are denoted with open lollipops. Bases artificially introduced during library preparation due to end repair or A-tailing are colored red. Fragments 633 635 637 639 641 643 645 depicted in the RRBS library prep start and end with the *Msp1* digest sites (CCGG) flanking the initial piece of genomic DNA. (B) Read pileups after mapping RRBS and WGBS libraries to a reference genome (red asterisks mark *Msp1* digestion sites). In contrast to data from WGBS, reads from RRBS libraries cover a small fraction of the genome. Further, because genomic DNA is fragmented with *Msp1* and then size selected (usually for fragments ~150-300bp in length), all fragments retained during the library prep should start and end with an *Msp1* recognition site and the pool of fragment will be enriched for CpG sites. Sequencing reads that are shorter than the original fragment length will localize to the *Msp1* recognition site associated with either the 5’ or 3’ end of the original fragment. (C) Pie charts showing the fraction of all CpG sites in the human genome covered by an RRBS experiment versus a WGBS experiment with similar total read depths (20 million reads generated *in silico*). Bar charts show, for each method, the proportion of measured CpG sites that fall in gene bodies (between the TSS and the TES), promoters (2 kb upstream of the TSS), CpG islands, and regions far from genes (>100 kb from any annotated TSS or TES).

High-throughput bisulfite sequencing data have a number of unique properties that influence both study design and data analysis. First, the raw data are binomially distributed count data, in which both the number of methylated reads (unconverted “C” bases) and the total read depth (number of methylated “C” bases plus unmethylated “T” bases) at each site contain useful information. For example, a site where 5 of 10 reads are methylated and a site where 50 of 100 reads are methylated both have estimated methylation levels of 50%. However, confidence in the methylation level estimate is higher for the second site, where total read depth is much greater. Information about relative confidence can be retained by modeling the raw count data rather than transforming counts to proportions or percentages, and several software packages now implement beta-binomial or binomial mixed effects models that do so14–16,30 (Box 1). These approaches provide a more powerful alternative to tests that assume continuously varying percentages or proportions (e.g., t-tests, Mann-Whitney U tests, linear models). They also control for count overdispersion, a known property of bisulfite sequencing data that violates the assumptions of commonly used, but false positive-prone^30^, binomial models.

Retaining count/read depth information during analysis also relates to a second property of bisulfite sequencing data: often, some samples will have low read depth or missing data at a CpG site where other samples have much higher read depth (especially in RRBS data sets, where read coverage is affected by the sample-specific efficiency and specificity of the restriction enzyme digest: Figure 1, Figure S1). Unlike RNA-seq data sets where read depth variation within a sample captures biological information (i.e., once normalized, lower read counts indicate lower expression levels), variance in read depth across sites in bisulfite sequencing data sets is purely technical and tells us nothing about biological variation in DNA methylation levels at a site. Both read depth and effective sample size will thus vary across sites in the same data set, and will often do so systematically across different regions of the genome (e.g., near genes or in intergenic regions, which differ in CpG content: Figure S1).

Finally, the efficacy of the bisulfite conversion step can vary across samples or groups of 116 samples prepared together, creating global batch effects. Though conversion efficiency is typically high (>98% of unmethylated cytosines converted to thymine^31–34^), small differences in conversion efficiency can have significant effects on genome-wide estimates of DNA methylation levels. In particular, samples with low conversion efficiencies will tend to have systematically upwardly biased estimates of DNA methylation levels relative to samples with higher conversion efficiencies, because fewer unmethylated Cs were converted to Ts. Thus, *sample-specific* bisulfite conversion rates should be directly estimated and taken into account in downstream analyses (e.g., by including bisulfite conversion rate as a model covariate). Estimates of sample-specific conversion rates can be obtained by spiking in a small amount of unmethylated DNA during library construction (lambda phage DNA is commonly used), mapping 126 the resulting reads to the appropriate genome (e.g., the lambda phage genome), and estimating the proportion of unmethylated cytosines in the spike-in DNA sample that failed to convert^31–34^. For data sets that do not include spike-ins, bisulfite conversion rates can be estimated using the 129 conversion rate of CHH and CHG sites in species or cell types in which CHH and CHG methylation is rare27,35,36. However, this approach is less ideal because it cannot differentiate between unmethylated cytosines that failed to convert because of technical reasons, and 132 methylated cytosines that truly occur in a non-CpG context.

## Effect sizes in ecological and evolutionary studies

A primary determinant of power in differential methylation analysis is the distribution of true effect sizes (i.e., the magnitude of the effect of the predictor variable of interest on DNA methylation levels). However, it is not obvious what the distributions of effect sizes for questions of ecological and evolutionary interest are likely to be. While effect size distributions and power analyses have been published for human disease case-control studies^18,20^, comparable 140 information is not readily available for most other settings. Small or moderate epigenetic changes may still impact gene expression levels and consequently be of interest^37–39^; however, they will require larger sample sizes to detect.

To aid researchers in choosing appropriate sample sizes, we estimated effect sizes in data sets from plants, hymenopteran insects, and mammals that address a range of ecological and evolutionary questions, including: (i) developmental and demographic effects (eusocial insect caste differentiation^35^; age^31^); (ii) ecological effects (resource availability, including both large differences^40^ and more modest ones^31^); (iii) genetic effects (*cis*-acting methylation quantitative trait loci^41^); and (iv) species differences^42,43^ (Table 1). For comparison, we also include a data set contrasting cancer cells with normal tissue from the same donors^21^, which produces some of the largest effect sizes for differential methylation observed to date.

**Table 1.** RRBS and WGBS data sets reanalyzed in this study

| Species | Predictor of interest | Contrast | Method | Citation |
| --- | --- | --- | --- | --- |
| Dog (*Canis lupus familiaris*) and wolf (*Canis lupus*) | Species differences | dog versus wolf | RRBS | 43 |
| Human (*Homo sapiens*) | Gestational famine during the Dutch Hunger Winter | famine-exposed versus same sex unexposed sibling | RRBS | 40 |
| Yellow baboon (*Papio cynocephalus*) | Age | continuous age values | RRBS | 31 |
| Yellow baboon | Resource base | wild-feeding versus human refuse-supplemented | RRBS | 31 |
| Clonal raider ant (*Cerapachys biroi*) | Caste | reproductive phase versus brood care phase | WGBS | 35 |
| Human | Cancer status | normal versus colorectal tumor samples (paired) | WGBS | 21 |
| Human, orangutan (*Pongo abelii*), gorilla (*Gorilla gorilla*), and chimpanzee (*Pan troglodytes*) | Species differences | human versus other great apes | WGBS | 42 |
| Mouse ear cress (*Arabidopsis thaliana*) | Local genetic variation | nearby (putatively *cis*-acting) genotype | WGBS | 41 |

We first reanalyzed each data set using a uniform analysis pipeline (Supplementary Materials) and estimated two measures of effect size: (i) the mean difference in methylation levels between groups of samples, for binary comparisons (Figure 2A) and (ii) the proportion of variance explained by the variable of interest (Figure S2). This analysis provides an empirical picture of how effect size distributions vary across study types. For example, local genetic variants tend to have large effects on DNA methylation levels, while environmental effects are consistently more modest (Figure 2A; Figure S2). To understand how these differences impact power, we simulated bisulfite sequencing data sets across a range of typical effect sizes and estimated the sample size required to identify differentially methylated sites in each case. All simulations presented in the main text assume that 10% of the sites in each dataset are true positives, but results from parallel analyses with varying proportions of true positives are shown in Figure S3.

**Figure 2.**
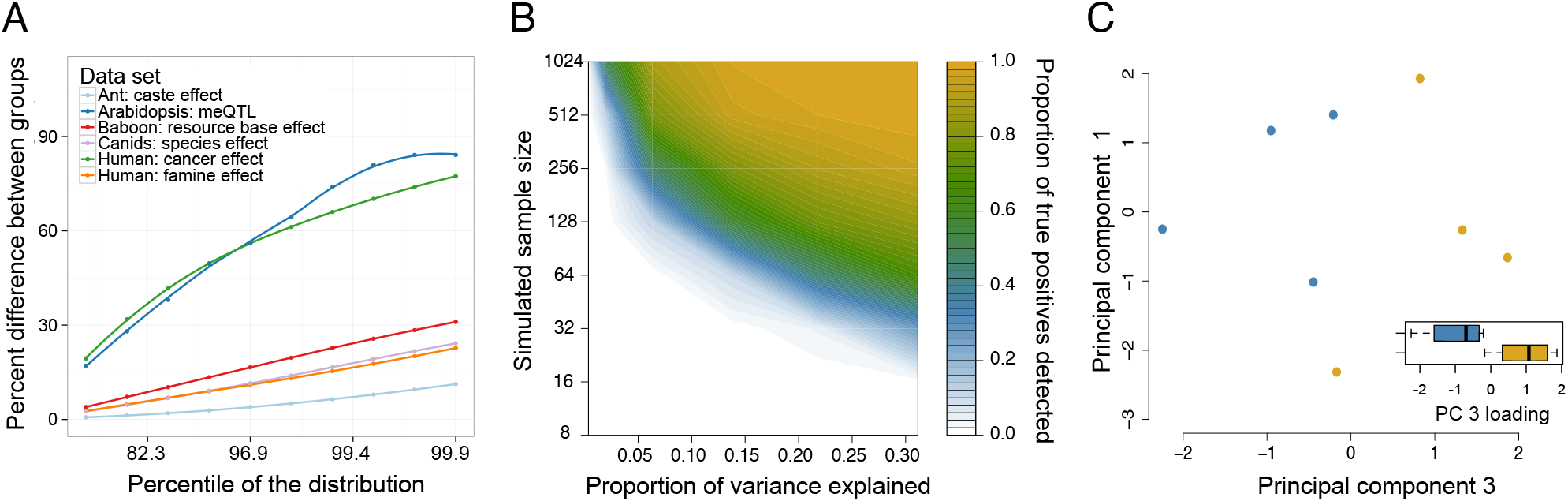
Estimates of effect sizes and their impact on the power of differential methylation analysis. (A) The maximum percent difference in mean DNA methylation levels between two sample groups (y-axis), for selected percentiles of sites (x-axis, ranked from smallest to largest percent difference) in reanalyzed data sets (Table 1) with binary predictor variables. Mean differences are based on raw values, without correction for covariates that may be imperfectly balanced between sample groups (e.g., age of study subjects in the baboon data set). We focus on the largest percentiles here because those effects are most likely to be detected or of interest to most investigators. (B) Power to detect differentially methylated sites at a 5% FDR in simulated RRBS datasets. Power increases as a function of the simulated sample size (y-axis; note that sample size is plotted on a log scale) and the magnitude of the effect of interest on DNA methylation levels (x-axis, represented as the proportion of variance in DNA methylation levels explained by the predictor variable). (C) In a small dataset (n=8), site-by-site analyses are too underpowered to detect differential methylation between clonal raider ants in the reproductive versus brood care phases. However, principal components analysis clearly separates samples by caste (t-test for PC 3, which explains 21.7% of the overall variance: p = 0.022; inset compares PC 3 loadings for reproductive versus brood care samples).

Our simulation results suggest that answering many ecological and evolutionary questions will require sample sizes that exceed those used in most current studies (Figure 2B; Table S1). For example, to identify sites where the predictor variable explains 15% of the variance in DNA methylation levels (a mean difference between sample groups of 13-14% in our simulations) with 50% power requires an estimated 125 samples (250 samples for 80% power and 500 samples for 95% power). To accommodate the costs of larger sample sizes, we recommend reducing per sample read depth or choosing a reduced representation or capture-based approach rather than WGBS. However, we strongly recommend against pooling DNA samples from multiple individuals into a single library, as this approach reduces power by collapsing the number of biological replicates available for analysis. Global analysis approaches that test for patterns in an entire data set, such as principal components analysis (PCA) or hierarchical clustering, may also be helpful when a data set is too power-limited to compensate for the large multiple hypothesis testing burden incurred in site-by-site analyses. This approach is particularly useful when a predictor variable is associated with small changes in DNA methylation levels at any given locus, but such changes are common genome-wide.

For example, in two published data sets (focused on the epigenetic effects of dominance rank in rhesus macaques and caste differences in clonal raider ants^32,35^), bisulfite sequencing sample sizes were very small. The macaque study (n=3 high-ranking versus n=3 low-ranking animals) did not attempt site-by-site analysis, while the raider ant study (n=4 pools of reproductive phase ants versus n=4 pools of brood care phase ants) found no evidence for caste effects on DNA methylation using site-by-site paired t-tests. As shown in Figure 2B (see also Figure S4), this result could have stemmed from low power. In support of this possibility, global analysis separates the sample groups of interest in both data sets. Specifically, the macaque study reported that hierarchical clustering distinguishes between high-ranking (n=3) 187 and low-ranking (n=3) individuals, with increased separation when focusing on CpG sites near genes differentially expressed with rank^32^. Similarly, when we re-analyzed the clonal raider ant data using principal components analysis, we found that a principal components analysis of CpG sites cleanly separates reproductive and brood care individuals, particularly along principal component 3 (t-test for separation along PC 3: p=0.022; Figure 2C). Together, these results 192 emphasize the potential utility of global analysis approaches in small studies.

## Kinship and population structure

Ecological and evolutionary studies often focus on natural populations that contain 196 related individuals or complex population structure. Accounting for these sources of variance is important because DNA methylation levels are often heritable^41,44–46^. In humans, where genetic 198 effects on DNA methylation have been best studied, genotype-DNA methylation associations have now been reported for tens of thousands of CpG sites^33,47–49^, with average heritability levels of 18%-20% in whole blood^45,46^. As a result, more closely related individuals will tend to exhibit more similar DNA methylation patterns than unrelated individuals. Analyses that do not take genetic relationships into account can therefore produce spurious associations if the predictor of interest also covaries with kinship or ancestry. For example, samples are often 204 collected along transects where climatic variables (e.g., temperature, altitude, rainfall) covary with genetic structure^41,50^. Genetic effects on DNA methylation could thus masquerade as climatic effects if genetic sources of variance are not also modeled.

Fortunately, this problem is structurally parallel to problems that have already been addressed in genotype-phenotype association studies, phylogenetic comparative analyses, and research on other functional genomic traits. The most straightforward solution is to use mixed effects models, which can incorporate a matrix of pairwise kinship or shared ancestry estimates to account for genetic similarity (Box 1). Specifically, this matrix is treated as the variance-covariance matrix for the heritable (genetic) component of a random effect variable (the environmental component is usually assumed to be independent across samples, so its variance-covariance is given by the identity matrix). The kinship matrix thus contributes to the predicted value of a heritable response variable, but does not affect the value of nonheritable response variables. Notably, while most approaches for controlling for relatedness implement linear mixed models that are only appropriate for continuous response variables^51–53^, recently developed binomial mixed models can be used to achieve the same task using count data^30^ (Box 1). These approaches avoid the need for transforming bisulfite sequencing data from counts to proportions or ratios, thus preserving information about sequencing depth for each site-sample combination. Additionally, recent tools for calling SNP genotypes directly from bisulfite sequencing reads (e.g., BisSNP^54^ and BS-SNPer^55^) can help with constructing kinship/relatedness matrices, although not without error (Box 2).

## Taxonomic differences in DNA methylation patterns

Most research on DNA methylation to date has focused on humans and a handful of model systems. However, ecologists and evolutionary biologists study a much broader range of species, and patterns of DNA methylation can vary dramatically among taxa^24,25^. These differences, too, can impact power and analysis strategies for bisulfite sequencing studies. They also mean that patterns typical of one taxonomic group cannot necessarily be extrapolated to others.

To provide some intuition about how the distribution of DNA methylation levels vary across taxa, we synthesized data from published studies of flowering plants, hymenopteran insects, canids, humans, and non-human primates (Table 1). We estimated the mean and variance of DNA methylation level for each CpG site in each data set (Figure 3A-B; Figure S5), and used these values to simulate new data sets for power analyses (Supplementary Materials). We were particularly interested in understanding the impact of variance on power because it is unlikely that a predictor variable of interest will significantly explain variation in DNA methylation levels at a locus where there is little variation to begin with. Importantly, the degree to which genomes are composed of relatively monomorphic (low variance) versus high variance sites systematically varies due to both taxon and sequencing strategy (Figure 3A-B, Figure S5).

**Figure 3.**
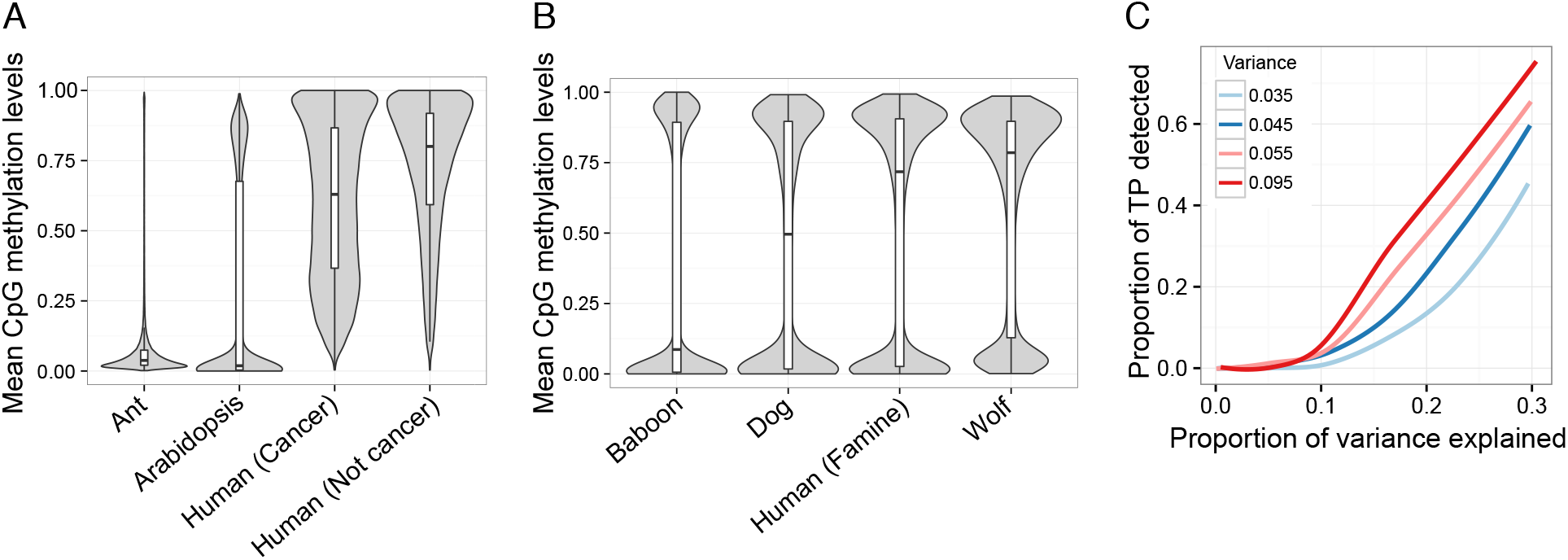
Properties of CpG methylation levels vary across data sets and influence power. For each (A) WGBS and (B) RRBS data set, we plotted the distribution of mean DNA methylation levels at each CpG site with a median coverage >10x across all samples in the study. (C) Power to detect differentially methylated sites (at a 5% FDR) in simulated RRBS datasets. The proportion of simulated true positives (TP) detected is plotted on the y-axis. Power increases as a function of the simulated effect size (represented as the proportion of variance explained; x-axis) and the variance in DNA methylation levels (colors). For all simulations, mean DNA methylation levels were held constant. The levels of variance in DNA methylation levels explored here (0.035, 0.045, 0.055, and 0.095) represent common values observed in real bisulfite sequencing data sets (Figure S5).

Our simulations suggest that, all else being equal, power to detect differential methylation in bisulfite sequencing data is limited by variance. Specifically, for any given sample size with a fixed mean DNA methylation level, power increases as a function of the underlying variance in DNA methylation levels (Figure 3C). These results suggest that analyses of low variance genomes, such as those typical of hymenopteran insects, may require larger sample sizes to detect a given effect than analyses of more variable systems, such as plants or mammals. An alternative, a less expensive approach is simply to filter out low variance sites prior to data analysis. Notably, such filtering will also affect the relative representation of sites in genes, promoters, CpG islands, and other functional compartments of the genome, because some of these compartments are consistently more variable than others (Figure S5).

In the current literature, differences in the genome-wide distribution of DNA methylation levels across taxa have led to taxon-biased analysis approaches. For example, in hymenopteran insects (where most of the genome is hypomethylated), several studies^35,56,57^ have used a binomial test to classify sites into ‘unmethylated’ or ‘methylated’ categories (i.e., all sites that do not pass a given significance threshold are considered ‘unmethylated’). Our simulations (Supplementary Materials) suggest that this approach not only loses information about quantitative variation, but is also sensitive to technical aspects of the data, such as sequencing depth. For example, using a binomial test approach, a site with an observed methylation level of 15% would be considered ‘unmethylated’ at a read depth of 20x, but ‘methylated’ at a read depth of 26x (Figure S6). This problem likely accounts for the report of high rates of ‘sample-specific DNA methylation’ (where a site is methylated in one sample, but unmethylated in all other samples) in one recent study^35^. Indeed, our re-analysis of the same data shows that 77% of putative sample-specific sites can be more parsimoniously explained by greater read depth in the “outlier” sample (Figure S6). Such problems can be readily avoided by not binarizing DNA methylation levels, which are intrinsically continuous traits, and by using count-based models that account for variation in sequencing depth^14–16,30^.

## Cell type heterogeneity

Epigenetic patterns vary substantially across cell types, contributing to differences in gene expression and biological function among different tissues^58^. In some settings, purified cell types can be isolated (e.g., via fluorescent-activated cell sorting). However, this approach is usually not feasible for biologists working under field conditions or with non-model systems (for which antibodies to cell type-specific markers are often unavailable). Consequently, most ecological and evolutionary studies have generated bisulfite sequencing data from heterogeneous samples, such as whole blood, whole organs, or even whole organisms. Because many variables influence both DNA methylation levels and cell type composition, putatively differentially methylated sites could, in some cases, be more parsimoniously explained by variation in cell type proportions rather than a direct effect of the variable of interest on DNA methylation^59^.

If isolating purified cell types is not an option, several alternative approaches can be used to address cell type heterogeneity. The best option is to directly estimate the relative proportion of the primary cell types in each sample using cell staining (e.g., Giemsa or Wright-Giemsa) or flow cytometry techniques. These estimates, or a composite measure of multiple estimates (e.g., the first several principal components of variation in cell type proportions) can be incorporated as covariates in downstream analyses. If no measures of cell type heterogeneity are available for the samples of interest, a second option is to use epigenomic profiles from sorted cells^36,60^ to predict the composition of mixed samples (a process known as ‘deconvolution’^59,61^). However, deconvolution estimates may introduce additional error, especially for cell types that occur at low frequency. A third option is to use data from sorted cell populations to understand the degree to which cell type composition could confound the conclusions of a study. If sites that are differentially methylated with respect to the predictor of interest also tend to be differentially methylated by cell type, the analysis may be confounded^6,31^. However, if the between-sample compositional differences that would be required to produce the observed levels of differential methylation are not biologically plausible, tissue heterogeneity is unlikely to completely explain observed differentially methylated sites^6^.

Finally, if data from sorted cell populations are unavailable, researchers can apply methods that account for cell type heterogeneity without the need for reference information^62–64^. However, caution is warranted, as some methods make implicit assumptions that may be violated in a given data set. For example, the program FaST-LMM-EWASher controls for cell type heterogeneity by (i) subsetting the data set to focus on the sites most strongly associated with a predictor variable of interest, and then (ii) calculating pairwise covariance between samples using only these sites. The resulting covariance matrix is included as a proxy for covariance in cell type composition in a mixed effects model^62^. However, FaST-LMM-EWASher makes two important assumptions: that most apparent cases of differential methylation are driven by cell type composition effects, and that true positive associations are therefore both rare and of large effect. These assumptions may hold in some studies, but when violated, this approach can substantially reduce power. For example, an analysis of resource base effects in baboon whole blood identified an association with DNA methylation levels at 1014 sites, after ruling out tissue heterogeneity confounds based on blood smear counts and comparisons against purified cell populations^31^. In comparison, FaST-LMM-EWASher detected a single differentially methylated site in the same data set. Alternative programs that account for cell type heterogeneity while making fewer assumptions (e.g., RefFreeEwas^64^ or SVA^63^) may thus be more appropriate. However, we caution that while such approaches can help control for variance due to cell type heterogeneity, none can overcome systematic *confounding* between cell type composition and a predictor of interest.

## Conclusions and tools

Like most other genomic technologies, high-throughput bisulfite sequencing approaches were first optimized in research contexts that afford a high degree of control (e.g., experimental case-control studies in model systems) and in systems that boast extensive genomic resources (e.g., humans). However, for ecologists and evolutionary biologists, these approaches often become most exciting when they can be extended to a much more diverse set of species and populations—even if these extensions come with complications. We believe that the biological insights to be gained from studies of DNA methylation in diverse taxa have substantial potential. However, maximizing the yield from these studies will require careful consideration of taxon-specific characteristics, the use of analysis methods appropriate to a data set’s structure, and realistic assessments of power. In particular, our results reveal that, with sample sizes that are currently feasible for many ecologists and evolutionary biologists, differential methylation analyses will tend to be moderately or lowly powered. Such studies may still have the potential to reveal interesting and important biology. However, researchers should be aware that they are likely to detect only the largest effect sizes (as is also true for other types of genomic analysis^65^), and should consider this bias when drawing biological conclusions.

Finally, to help quantify how sample size, effect size, population structure, and modeling approach affect bisulfite sequencing data analysis, we have developed an R Shiny application to perform power analyses like those presented here. This app allows bisulfite sequencing data to be simulated with user-specified properties, is coupled with a set of statistical analysis options to evaluate study power, and outputs the simulated count data for maximal flexibility. The app is freely available at www.tung-lab.org/protocols-and-software.html.

## Acknowledgements

We thank Kasper Hansen and Irene Hernando-Herraez for providing processed file formats from their previously published work. We also thank Noah Snyder-Mackler, Luis Barreiro, and Xiang Zhou for helpful comments and suggestions, Mine Cetinkaya-Rundel for coding suggestions on the R Shiny app, and the Baylor College of Medicine Human Genome Sequencing Center for access to the current version of the baboon genome assembly (*Panu 2.0*). This work was supported by NIH R21-AG049936 and 1R01GM102562 to JT, NSF BCS-1455808 to JT and AJL PAPD is supported by NIH K12GM000678 from the Training, Workforce Development & Diversity division of the National Institute of General Medical Sciences.

## Data accessibility

Summaries of data availability and accession numbers for previously published data sets are provided in Table S1. Our BisSNP analyses utilized publicly available SNP calls for *Arabidopsis* accessions downloaded from the 1001 Genomes Project (http://1001genomes.org/data/GMI-MPI/releases/v3.1/). An R Shiny app for simulating bisulfite sequencing data and performing power analysis is available at www.tung-lab.org/protocols-and-358software.html.

## Author contributions

AJL and JT conceived the study; AJL, TPV, and PAPD analyzed previously published and simulated data; TPV wrote the R Shiny app; and AJL and JT wrote the manuscript, with input from all co-authors. All authors gave final approval for publication.

### Box 1. Modeling approaches for bisulfite sequencing data

*Binomial regression.* A **binomial distribution** intuitively describes bisulfite sequencing data generated for a given sample, *i*, at a given site: the number of methylated counts *(m)* represents the number of ‘successes’ in an experiment with *t* trials and *p* probability of success. Here, *t* translates to the total read depth and *p* to the (unobserved) true methylation level.

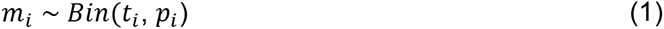

However, bisulfite sequencing data are overdispersed (i.e., show greater variance than expected) relative to binomial expectations. Thus, using a **binomial regression** to model bisulfite sequence data can result in an extremely high rate of false positives and is not recommended^14,30^.

*Beta binomial regression.* To account for overdispersion, **beta binomial regressions** have been proposed for bisulfite sequencing data^14–16^. Here, the parameter *p_i_* from the binomial setting (equation 1) is itself treated as a random variable that follows a two-parameter beta distribution.

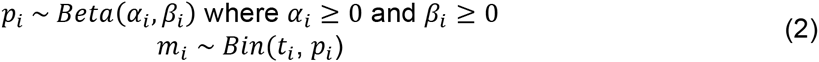

The beta distribution is then re-parameterized as a beta binomial with parameters *t_i_*, *π_i_* (equal to *α_i_*/(*α_i_, + β_i_*)), and *γ* to capture overdispersion.

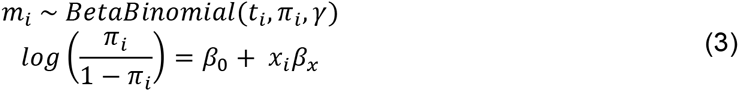

Here, *π_i_* is the analog of the binomial probability of success (*p_i_*) and can be interpreted as the underlying true methylation level (note that the binomial distribution is a special case of the beta binomial distribution when *γ*=0). *π_i_* is passed through a logit link function in order to transform probability values (which are bounded between 0 and 1) to a continuous space for linear modeling. Transformed values are modeled as a function of an intercept (*β_o_*), the predictor variable of interest (*x_i_*), and its effect size (*β_x_*).

*Linear mixed effects models.* While beta-binomial regressions have become a popular tool for modeling bisulfite sequencing data, these models are not appropriate for data sets that contain related individuals or population structure. Such data sets require approaches that can account for genetic covariance (i.e., nonindependence) among samples, such as **linear mixed effects models**.

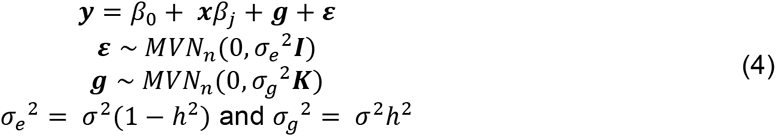

Here, *y* is a vector of continuously distributed methylation levels (obtained by normalizing *m/t*) and *g* is a vector of random effects with a covariance structure determined by the genetic relatedness among individuals in the sample (described by ***K***, a user-defined n x n pairwise relatedness matrix) and the heritability of the DNA methylation trait (*h^2^*, which can be decomposed into its genetic and environmental components). *I* is an n x n identity matrix.

*Binomial mixed effects models.* Linear mixed models are flexible and fast, but discard information about total read depth when counts are normalized. **Binomial mixed effects models** overcome this constraint by controlling for genetic covariance while modeling raw counts.

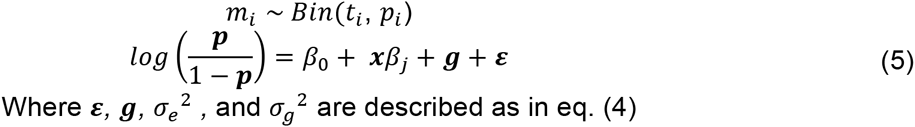

This model essentially combines the linear mixed model with the beta binomial regression. The variable *p* now reflects the vector of true methylation levels for all samples and is passed through a logit link function for linear modeling. The genetic covariance, as well as the overdispersion, is captured by the random effects component.

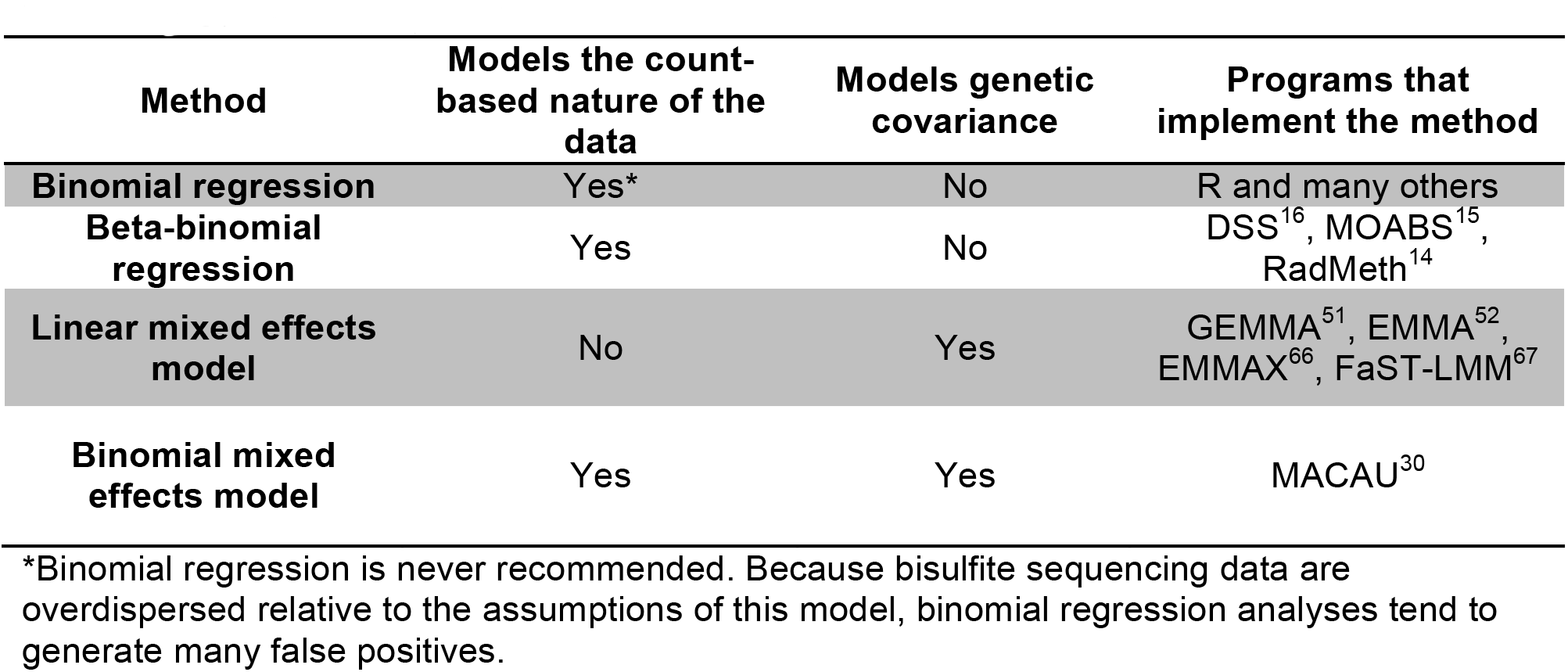
Summary of model properties

### Box 2. Calling genotypes from bisulfite sequencing data

Like other high-throughput sequencing assays^68,69^, bisulfite sequencing studies generate sequencing reads that contain information about genetic variation. Calling variants or genotypes from these data may be of interest for detecting genetic effects on DNA methylation levels (i.e., methylation quantitative trait loci, or meQTL), verifying sample identity, or controlling for genetic relatedness in downstream analyses. However, typical SNP-calling algorithms are not well suited to bisulfite sequencing data because the C to T conversion obscures true C/T polymorphisms. Several recently developed software packages attempt to overcome these challenges^54,55^. To assess the performance of one such program, BisSNP^54^, we analyzed a whole genome bisulfite sequencing data set for 29 *Arabidopsis thaliana* accessions^41^ where SNP calls were also available from whole genome sequencing through the 1001 Genomes Project and, for a subset of these individuals (n=25), genotype array data^70^.

Using BisSNP under default recommendations (Supplementary Materials), we identified 235,338 biallelic variable sites. This set was highly skewed to transitions (n=234,512 transitions, 99.65% of all called sites). Only 45% (n=106,925) of variants called using BisSNP represent putatively ‘true’ variants that were also identified in the 1001 Genomes resequencing data, but transversions were much more likely to be ‘true’ variants than transitions (90.3% compared to 45.3%). More stringent variant call filtering (variant quality ≥50 rather than ≥30) increased the proportion of likely true variants to 50.3%, but at the cost of retaining only 4.7% of the original sites. However, for previously identified variants in the BisSNP call set, BisSNP genotype calls and genotype array data agreed 87.5% of the time, with transversions agreeing more often than transitions (93.1% compared to 87.4%). Thus, BisSNP appears to provide relatively high-quality genotyping information for known variants.

However, our analyses do suggest that BisSNP genotypes provide a reliable way to verify sample identity and capture population structure. Using the set of biallelic SNPs that were identified by BisSNP, the 1001 Genomes Project, and the array data (n=3,553 SNPs overlapped between all 3 methods for n=25 accessions), a neighbor joining tree^71^ clearly clusters samples by accession. The single exception was a WGBS sample that may be mislabeled, as the BisSNP calls clustered separately from the resequencing and array genotype calls for this accession. Further, the pairwise genetic covariance matrix generated from BisSNP calls was highly consistent with the genetic covariance matrix generated from whole genome resequencing data (Mantel test r = 0.873, p < 10^-6^). Perhaps more importantly, the differences we did detect had marginal effects on differential DNA methylation analysis. Specifically, when we analyzed possible methylation quantitative trait loci (meQTL) in the *Arabidopsis* data set (Supplementary Materials), meQTL effect sizes were highly consistent between analyses using BisSNP calls to estimate population structure and analyses using whole genome resequencing data (Spearman’s rho=0.925, p<10^-15^).

